# A single-cell genetic colocalization test improves power and resolves disease-mediating cell types

**DOI:** 10.1101/2025.10.10.681685

**Authors:** Jonathan Mitchel, Sumaiya Nazeen, Xinyuan Wang, Praveen Kumar Patnaik, Autumn Morrow, Habib Nasir, Ronya Strom, Dylan Ritter, Lorenz Studer, Sung Chun, Chris Cotsapas, Vikram Khurana, Peter V. Kharchenko, Shamil R. Sunyaev

## Abstract

Statistical colocalization testing methods can determine if the same single-nucleotide polymorphism (SNP) underlies both a genome-wide association study (GWAS) locus as well as an expression quantitative trait (eQTL) locus. This can nominate potential mechanistic pathways from SNPs to genes to traits, while providing cell type or tissue context. Surprisingly, systematic colocalization testing with bulk-tissue eQTLs fails to link the majority of GWAS loci with gene expression changes. Mapping eQTLs with single-cell expression data has the potential to reveal the missing regulatory effects of GWAS variants. However, current pseudobulk cluster-based approaches may be underpowered when clustering accuracy is imperfect or with an incorrectly selected cluster resolution. To improve power of single-cell colocalization tests, we developed a cluster-free method, scJLIM. By modeling eQTL interactions with continuous cell states (e.g., principal components), scJLIM estimates eQTL significance and colocalization in individual cells. We benchmarked our method with simulated data, demonstrating improvements in power over pseudobulk methods. In our main applications, we used scJLIM to analyze blood and brain scRNA-seq datasets paired with autoimmune and neurological disease GWAS, respectively. We identified nearly twice as many total colocalizations compared with traditional pseudobulk analyses carried out within the major cell populations of these tissues. Aligning with a recent experimental study, we highlighted an example of the *ETS2* gene colocalizing with an inflammatory bowel disease GWAS locus in a subset of myeloid cells. For Parkinson’s disease (PD), our results pointed to *TRPV2* as a potential gene of interest, corroborated by transcriptional changes in both post-mortem PD brains and iPSC-derived neuronal models of alpha-synucleinopathy.

## Introduction

Genome-wide association studies (GWAS) have unveiled many causal loci contributing to common diseases. However, it has been less successful than initially imagined at helping gain insight into the intricate cellular processes downstream of these pathogenic variants. It has been estimated that over 90% of trait-associated single nucleotide polymorphisms (SNPs) reside in noncoding sequences^1^ and that the majority of SNP heritability can be localized to non-coding regulatory regions marked, for example, by accessible chromatin (DNase I hypersensitivity sites)^2^. These studies and others have painted a picture of common variants impacting their downstream traits via gene regulatory mechanisms as opposed to changing protein structure. This gave rise to a natural, and seemingly the only existing, hypothesis that regulatory variants impact complex traits by modulating expression of relevant genes.

By generating RNA-sequencing datasets from many individuals paired with genotype data, it became possible to map expression quantitative trait loci (eQTLs) and test whether disease associated SNPs also modulate gene expression levels. Such evidence might suggest that the SNP’s causal effect on disease is mediated through this expression change, further providing evidence of the genes and tissues where that effect takes place. Since GWAS is unable to reveal the exact causal SNP due to linkage disequilibrium (LD), however, it is necessary to consider the full information from a given locus in what is referred to as a “colocalization test”. Colocalization methods indirectly infer whether the same SNP underlies both a disease trait and gene expression by taking LD into account when comparing SNP significance between traits. Each of the different colocalization methods make various assumptions, but in general, when colocalization exists for a given locus, we expect to see similar association patterns for SNPs in the locus between the disease and expression traits^3–6^.

Surprisingly, in studies of bulk tissue expression^6–9^, only 25-40% of GWAS peaks have been found to co-localize with any eQTL irrespectively of relevance of the regulated genes. Requiring that the downstream genes have strong biological evidence supporting their role in disease biology results in this fraction falling below 10%^10^. On the flip side, many eQTLs controlling expression of highly relevant genes in highly relevant tissues do not have any detectable effects on corresponding human phenotypes. The fraction of heritability of human phenotypes mediated by bulk gene expression has also been estimated at shockingly low 10%^11^. This poses a problem of “missing regulation”^10^ (in analogy of “missing heritability”).

There are several potential reasons why most GWAS fail to colocalize with eQTLs. These include low eQTL mapping power due to small sample sizes, context-dependent function of disease SNPs (e.g., if they are only active during development or in a stimulated state), or cell-type-specific effects that dampen/modulate the eQTL significance patterns observed in bulk tissue^12^. It seems unlikely that we lack power to identify eQTLs for important disease genes, as numerous highly significant eQTLs have been mapped for these genes. It is also uncertain to what degree of importance development or environmental context will resolve the missing regulation. One study conducted eQTL mapping in cultured fetal neurons and progenitors, identifying a handful of new colocalizations with neurological diseases not identified using adult bulk tissue^13^. Other studies have mapped eQTLs in *in vitro* stimulated cells^14^ or in clinical samples to test for interactions with phenotypic covariates^15^ (e.g., inflammation), identifying a handful of new colocalizations for immune-related traits. While promising, it appears that many non-coding GWAS loci remain unexplained by eQTLs.

Several studies have indicated the importance of moving to single-cell RNA-sequencing (scRNA-seq) data for resolving this question. Since non-coding GWAS variants likely alter transcription factor binding, their eQTL effects might differ by cell type. Early evidence for this came from a study by the GTEx consortium, which identified nearly twice as many colocalizations using cell type proportion interaction eQTLs (ieQTLs) compared to standard bulk eQTLs^16^. With technological advances in scRNA-seq sample multiplexing, it became feasible to generate scRNA-seq datasets from many individuals and map eQTLs directly with this modality^17^. One study conducted single-nucleus RNA-seq (snRNA-seq) on brain prefrontal cortex samples from postmortem, aged individuals and identified 40% more eQTLs not identified in matching bulk tissue, despite the bulk tissue having over twice the sample size^18^. A different study conducted scRNA-seq of T cells to identify eQTLs with effects that were dependent on the cell’s continuous-valued cell-state. These cell state-dependent eQTLs were enriched for overlap with GWAS variants over non-interacting eQTLs^19^. These studies point to single-cell data as a promising avenue for revealing some of the remaining missing regulation.

Current approaches to eQTL mapping with scRNA-seq data are typically conducted separately per cell type or subtype. One strategy is to collapse cells per cell type to a single sample for each individual, a process typically referred to as “pseudo-bulking”. Then, this transformed expression data can be used as input in standard bulk eQTL mapping^20^. Another strategy is to fit a mixed model with the individual cells as input, while including random effects per individual to account for the fact that cells from the same individuals tend to be more similar^20,21^. Other more complex models such as the cell-state-interaction eQTLs discussed above, use mixed models with interaction terms between genotype and some representation of cell states^22,19^. These representations are usually derived from principal component analysis (PCA) or other matrix decompositions applied to the scRNA-seq gene expression matrix. Thus, cell-state-interaction approaches make better use the rich heterogeneity to be found in scRNA-seq data and do not require making assumptions about the true cellular resolution at which an eQTL takes effect. They also avoid the use of cell clustering which can be inaccurate, unreproducible (especially at fine clustering resolutions), and misinterpreted given the use of select marker genes for annotation.

An ideal single-cell colocalization test would yield significance when colocalization exists in at least some of the cells tested. Therefore, the direct use of cell-state interaction significance as the eQTL statistics is often ineffective for colocalization testing. Using this strategy, a positive colocalization would indicate that the SNP underlying the GWAS is also a dynamic eQTL over the tested cell-state axis. However, this would fail if a subset of the cells harbored distinct eQTLs in the locus (non-colocalizing), as this would produce a significance pattern reflecting all dynamic eQTLs. It would also fail in cases where the true colocalizing cells require multiple PCs or axes for their separation. This approach also cannot pinpoint the specific cells that have the colocalizing eQTL.

To overcome these challenges while fully leveraging the rich cell-state information available, we developed scJLIM, the single-cell variant of our “Joint Likelihood Mapping” (JLIM) colocalization tool^6^. By broadly applying scJLIM to the largest single-cell immune and brain datasets, we uncovered roughly twice as many colocalizations compared to traditional pseudobulk methods. This highlights the importance of moving beyond arbitrarily defined clustering for eQTL mapping and interpreting GWAS loci. Overall, scJLIM advances our understanding of human diseases by revealing the genes, pathways, and cellular contexts through which GWAS signals exert their effects.

## Results

### scJLIM method overview

Several commonly used colocalization tools include COLOC^3^, JLIM^6^, fastENLOC^23^, and eCAVIAR^24^, among others. Each takes as input a set of GWAS and eQTL summary statistics and outputs either a p-value or posterior probability of colocalization. The novelty of scJLIM is not in the calculation of the colocalization statistic but instead in the eQTL mapping step. Uniquely, scJLIM estimates eQTL effects in individual cells conditional on each cell’s expression state, enabling the application of existing colocalization tools to individual cells. To do this, we first compute a low dimensional representation cell states using PCA on the gene expression matrix (Methods). Like with other colocalization tests, we also select a specific GWAS locus and gene to test (referred to as the “eGene”) (Figure 1, step 1). When a GWAS causal SNP is an eQTL only in a specific cell state, we expect to see large genotype-by-PC interaction effects on expression for PCs that separate out the specific cell state. Cells with that state will consequently have a total genotypic effect (i.e., marginal effect) and significance for SNPs in LD with GWAS causal SNP. To estimate these effects, scJLIM maps an eQTL for each SNP in a locus by using a linear mixed model with a set of genotype-by-PC interaction terms among other covariates (Figure 1, step 2) (Methods). With the fitted models, scJLIM then estimates the total genotypic effect, error, and significance of each SNP in each cell, given each cell’s unique location in PCA space.

**Figure 1.**
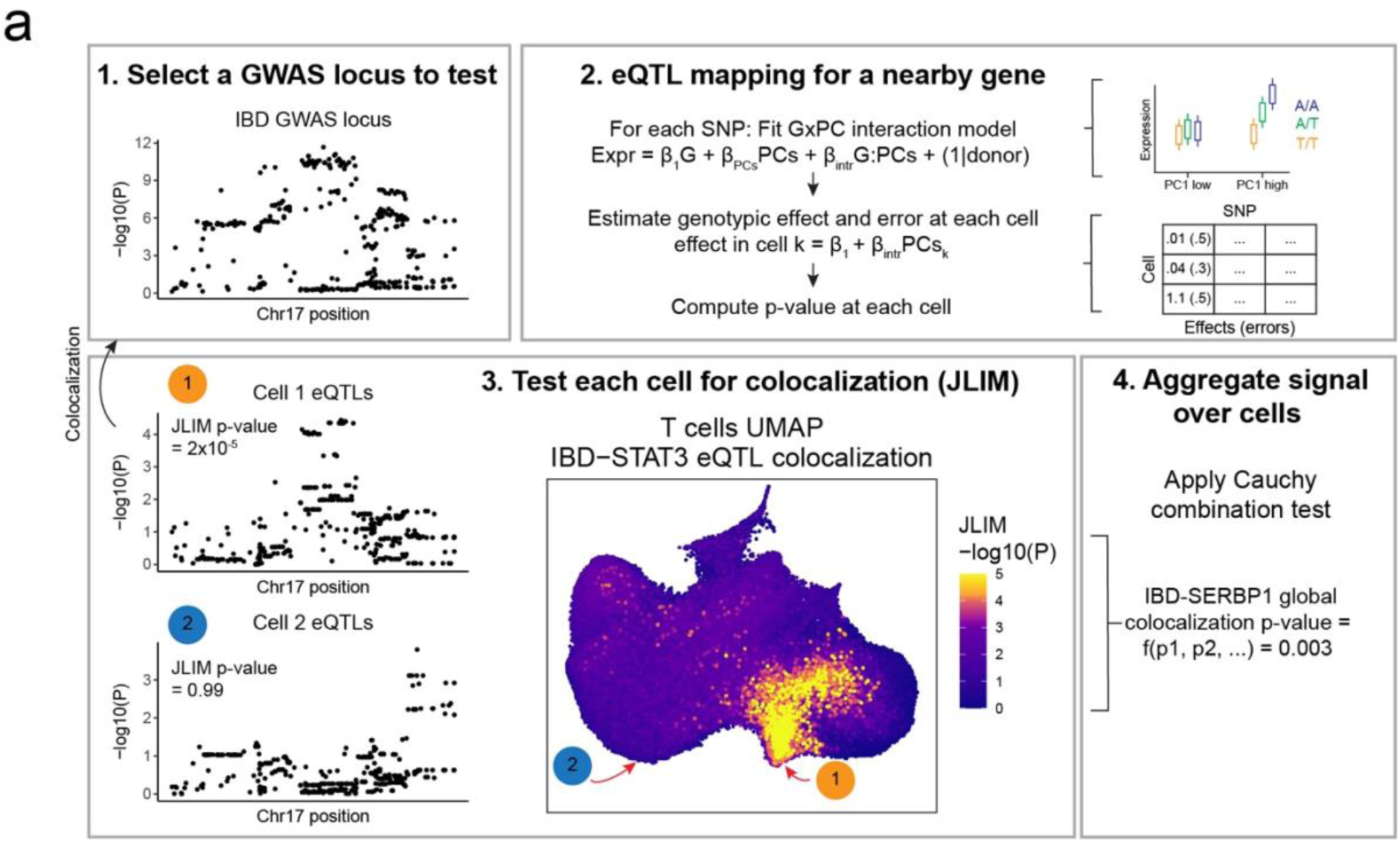
scJLIM method overview. a. After a GWAS locus is selected and paired with nearby genes for testing (step 1), scJLIM fits linear interaction mixed models for each SNP in the locus (step 2). Each of these includes a genotype term, other donor and cell-level covariates, PC covariates, and several genotype-by-PC interaction terms. Then, the fit model parameters are used to estimate the total genotypic effects and errors in individual cells conditional on their PC locations, yielding a Z statistic that we convert to a p-value. Each cell’s p-values over SNPs are used in a JLIM colocalization test with the corresponding GWAS locus (step 3). Finally, we compute a global p-value over all cells using the Cauchy combination test (step 4).

With estimates of eQTL significance across the cell state space, we can then apply standard colocalization testing methods to identify cell states where the significance profile matches that of a GWAS locus (Figure 1, step 3). For the colocalization testing component, we rely on the JLIM method^6^ as it provides a p-value which is necessary later on for multiple hypothesis test corrections. Briefly, JLIM iterates over SNPs in high LD with the lead GWAS SNP, computing a likelihood-based statistic that upweights SNPs with strong associations to both traits—unless more significant eQTLs exist outside of a specified LD window^6^. JLIM computes an empirical p-value from comparing the observed test statistic to a null distribution generated with null eQTLs. We then apply the Cauchy combination test^25^ to the set of JLIM p-values from all cells, yielding a single global colocalization p-value for the gene-locus pair, indicating whether there was significant colocalization among the tested cells (Figure 1, step 4).

### Testing and benchmarking scJLIM with simulated data

To construct simulated GWAS and scRNA-seq data, we first simulated genotype panels, using Hapgen2^26^ combined with LD information from the 1000 Genomes Project^27^ (Methods). We selected a single causal variant per locus and used these variants to create quantitative phenotypes for running a GWAS. For each set of GWAS results, we also generated 5 simulated scRNA-seq datasets each programmed with a cell-type-specific genotypic effect on expression for the corresponding GWAS causal SNP^28^ (Figure 2a left). We created two distinct positive scenarios. In the first scenario, the GWAS causal SNP had an eQTL effect in one cell population, with no other SNPs having programmed eQTL effects in any cell populations. In the second scenario, the GWAS causal SNP again had an eQTL in one cell population. However, in this scenario, the other cell populations contained programmed eQTLs only for SNPs that do not causally affect the GWAS trait. Both scenarios should yield significant global p-values, as both have programmed colocalization in at least one population of cells. We also used this simulation setup to confirm scJLIM’s ability to correctly nominate cell populations that contain the colocalizing eQTL, with an example shown in Figure 2b.

**Figure 2.**
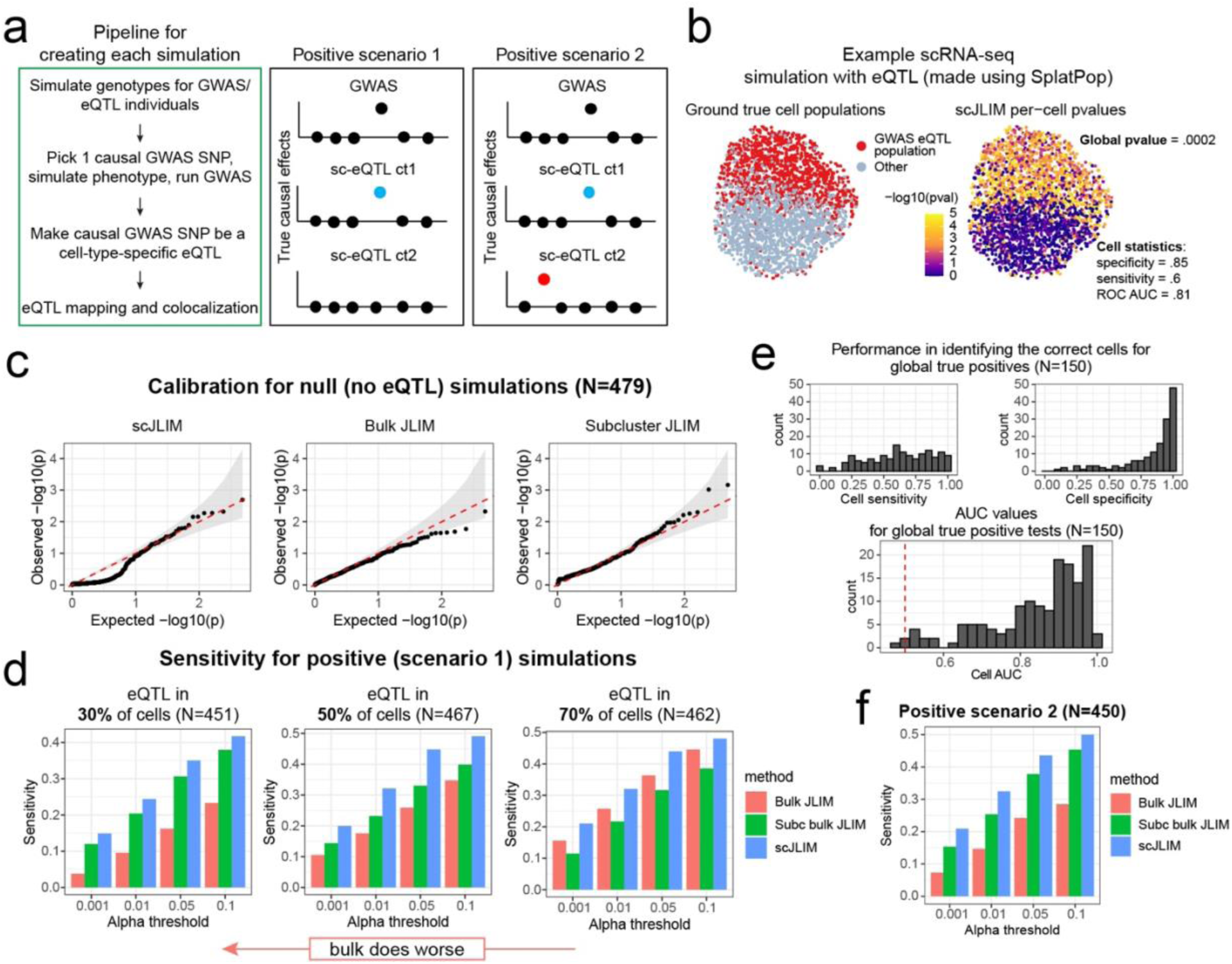
Benchmarking scJLIM with simulated GWAS and cell-type-specific eQTL data. a. Overview of the simulation generation pipeline (left) and two different positive scenarios designed (right). For each positive simulation scenario, we also generated matching null scenario that lack the eQTL corresponding to the causal GWAS SNP. b. An example of one simulated scRNA-seq dataset with a cell-type-dependent eQTL (left). Also shown are the per cell colocalization p-values (left). c. Calibration results for the null scenario of no eQTLs present with two cell types. We plotted the observed versus expected nominal global p-values. d. Global sensitivity results for positive simulation scenario 1 with two cell types. The cell population programmed to have the eQTL was made to be smaller than, the same size as, or larger than the population without the eQTL to demonstrate how relatively small populations are especially difficult to detect with bulk colocalization due to signal dilution. The arrows below these plots highlight the relative regimes where each method improves. e. Cell sensitivity, specificity, and ROC AUC for detecting the ground true cells that were given the eQTL for the same causal SNP as the GWAS data. This computed using the two-cell type, balanced-proportions, positive-scenario 1 simulations that yielded a global p-value less than 0.01. Significantly contributing cells were selected as those which pass our elimination-based procedure with the Cauchy combination test (Methods). f. Global sensitivity results for positive simulation scenario 2 with two cell types. The eQTL population was the same size as the non-eQTL population in these simulations.

Before testing the positive scenarios, we first evaluated the global p-value calibration of scJLIM using two-cell type simulations with no eQTLs present (Figure 2c). We also computed this metric with bulk JLIM (pseudobulked over all cells) as well as subcluster JLIM. The subcluster JLIM approach entails subclustering cells, pseudobulk eQTL mapping in each cluster, running JLIM per cluster, and meta-analyzing JLIM p-values over the clusters. These results demonstrate that scJLIM has comparable calibration to the other methods, especially in the lower alpha levels which are important for distinguishing true positive signals in real data. Results were similar for four-cell-type-simulations and a negative scenario 2 (GWAS causal SNP no longer an eQTL) (Supp. Figure 1a,b,c). As with the original JLIM method, we further show that the negative scenario 2 tests become more conservative as the eQTL effect becomes stronger and less difficult to discern from the GWAS signals (Supp. Figure 1b). We also note that scJLIM calibration becomes more conservative than the other methods at higher alpha levels. This is likely due to the Cauchy combination test itself, as its type I error rate is more accurate at the tails of the distribution^25^.

Next, we benchmarked sensitivity of each method in positive scenario 1 (Figure 2d). When the eQTL population was small (30% of all cells), we observed that scJLIM had double the sensitivity of bulk JLIM. Bulk JLIM approached the scJLIM sensitivity as the eQTL population increased in size. Bulk JLIM had reduced power with small eQTL cell populations because the inclusion of additional null cells increases noise without increasing the eQTL signal. Bulk JLIM frequently failed to yield significant colocalization when other eQTLs (distinct from causal GWAS SNP) were present in the second cell type (positive scenario 2), as this generates a locus-wide significance pattern that includes both eQTL signals (Figure 2f). This highlights the importance of disentangling such signals either through subclustering or with a cluster-free approach such as JLIM. We then tested how cluster accuracy impacted the sensitivity relative to the other methods. We noted that subcluster JLIM performed worse than bulk when clustering accuracy was low, and conversely, it matched the performance of scJLIM when run with ground-truth clustering (Supp. Figure 2b right) (Methods). This indicates how the performance of subcluster based colocalization is unstable in the face of imperfect clustering accuracy, whereas scJLIM is more robust to it with roughly two times higher sensitivity. We also note that in a more realistic situations with a more complicated cell type hierarchy, the relevant level of hierarchy (clustering granularity) is unknown. This creates an additional complication for the clustering-based approach.

For scJLIM global true positive tests (Cauchy p-value<0.01), we confirmed that our method could identify the subsets of cells that harbored the colocalizing eQTLs. We computed cell sensitivity and specificity (after selecting significant cells, Methods), and we calculated the receiver operating characteristic area under the curve (ROC AUC) with per-cell p-values (Figure 2e). Most importantly, specificity was very high, suggesting that scJLIM does a good job at not identifying the incorrect cell populations. Sensitivity ranged widely, but this is of less concern, as we nearly always had at least some cells of the correct population called significant. Results for the other scenarios were similar (Supp. Figure 2c,d). This demonstrates that scJLIM performs as intended when there is a positive signal present in a subset of cells.

### scJLIM uncovers many new colocalizations for autoimmune traits

We applied scJLIM to autoimmune disease datasets to identify subpopulations of immune cells that colocalize with autoimmune GWAS loci. We used the OneK1K scRNA-seq dataset of peripheral blood mononuclear cells (PBMCs) from 982 individuals, with paired SNP array genotype data needed to map eQTLs^29^. We applied standard preprocessing steps (Methods) and incorporated additional quality control data to remove doublets from this dataset^30^. We also ran batch correction using ComBat^31^, prior to computing PCA (Methods) within each of the previously annotated major immune cell categories (T cells, B cell, NK cells, and Myeloid cells). We downloaded GWAS datasets from 10 studies of autoimmune or immune-related traits^32–39^ and selected loci at least 100kb in size (Methods) that contained genome-wide significant SNPs with minor allele frequencies above 0.10. Each locus was paired with protein-coding genes within 500kb of the locus’ center for eQTL mapping. We applied scJLIM within each of the four major cell types to identify subpopulations that have eQTL colocalization with the GWAS loci. We also ran bulk JLIM after creating pseudobulk samples for individuals by aggregating their respective cells from the major cell type (Methods). Subcluster JLIM was run using pseudobulk data at the level of previously annotated subclusters and meta-analyzed over all subclusters per major cell type. This provided us with three sets of results for each major cell type (Supplementary Tables 1). In comparing these results, we first tabulated the number of unique loci per trait that yielded significance (Padj<.10) with each of the three methods (Figure 3a). These global p-values were adjusted using the Benjamini-Hochberg method to control the false discovery rate^40^. For 6 out of 10 of the GWAS datasets, scJLIM yielded nearly twice as many significant colocalizations as either bulk or subcluster JLIM. We further tabulated the number of significant gene-locus pairs in each trait-cell type combination tested and computed a summary meta-analysis p-value for each trait cell type pair (Figure 3b). Both bulk and scJLIM yielded significant colocalizations in multiple cell types for all traits except eczema (which only had one significant colocalization). The meta-analyzed p-values helped discern which cell types had the strongest signals per trait, for instance pointing to T cells for RA, B cells for PBC, B cells for SLE, and T cells in UC. The same trait-cell type links have also frequently been identified using other methods. For example, the S-LDSC method applied to bulk immune cell expression data previously identified heritability enrichment correspondingly in T cells for RA, in B cells for PBC, B cells for SLE, and T and NK cells for IBD^41^.

**Figure 3.**
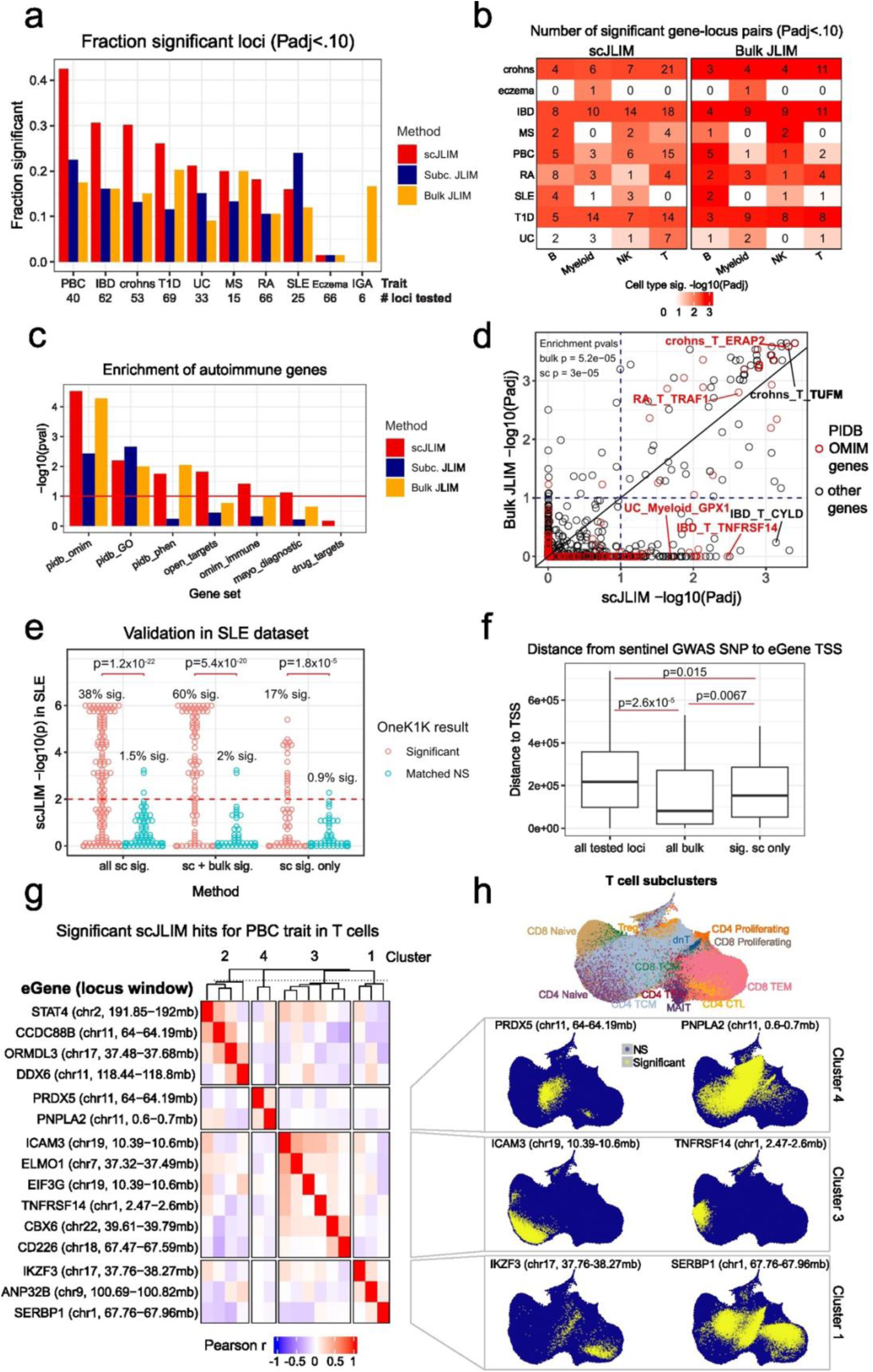
Application of scJLIM to the OneK1K scRNA-seq dataset and ten autoimmune disease GWAS datasets. a. Comparing the fraction of tested loci that are significant with either scJLIM, subcluster pseudobulk JLIM, or bulk JLIM. These results are aggregated over all four major cell types. The total number of tested loci are written in above each set of bars for a trait. b. The number of significant gene-locus pairs for each trait in each cell type, shown separately for scJLIM (left) and bulk JLIM (right). Color indicates p-values from meta-analyzing all tests per trait-cell type pair with the Cauchy combination test. c. We tested for enrichment of known autoimmune and immune genes among the significant eGenes by either scJLIM or bulk JLIM. These included the Pi autoimmune database disease, immune function, and immune phenotype genes (pidb_omim, pidb_GO, pidb_phen), as well as an autoimmune gene set from OpenTargets, OMIM immune-related disease genes, a Mayo clinical autoimmune gene diagnostic panel, and a list of autoimmune drug targets. Enrichment was computed with a hypergeometric test. d. Comparing scJLIM gene-locus pair global p-values versus bulk JLIM p-values. Highlighted in red are genes found in the pidb_omim gene set, which corresponds to the enrichment p-values found in the upper right as well as in panel (b). e. Validating the scJLIM hits from OneK1K in an SLE dataset. We show the -log10(p-values) in the SLE dataset for the hits which were originally scJLIM significant only or also significant with bulk JLIM. Each group is compared against a cell-type matched test for a locus which was nonsignificant in the OneK1K data. We tested whether the OneK1K significant hits were enriched (hypergeometric test) for having a significant hit in the SLE dataset compared to the OneK1K non-significant hits. f. For each significant gene-locus pair for each trait in each cell type, we calculated the distance from the sentinel GWAS SNP of that locus to the transcription start site of the eGene. These distances are shown separately for all tested gene-locus pairs, all bulk JLIM significant pairs, or the pairs which are significant only for scJLIM. Significance was calculated using two-sample Student’s t-tests (two-sided) on the natural log of the distances. g. Heatmap showing correlations between scJLIM significant (Padj<0.1) gene-locus pairs for the PBC trait in T cells. The correlations were calculated using the per-cell p-values in all T cells, which were transformed to Z-scores prior to calculating each correlation. Pairs of results were clustered using hierarchical clustering and cut to select four clusters. h. Showing UMAP plots highlighting which cells contributed to global significance for select gene-locus pairs in (g). Significantly contributing cells were selected using the scJLIM per-cell p-values and include only those which pass our elimination-based procedure with the Cauchy combination test (Methods).

We tested whether the eGenes identified by each method were enriched for genes with known immune functionality and autoimmune disease relevance (Figure 3c). These gene sets were derived from a variety of sources including autoimmune drug targets, OMIM, Gene Ontology, OpenTargets^42^, and others^43,44^. The scJLIM results were enriched in these gene sets at least as much as, if not more than the results from the other methods. We also show a comparison of the scJLIM versus bulk JLIM adjusted p-values highlighting the genes from one such gene set (Figure 3d). In general, the most significant bulk JLIM results were nearly always also identified with scJLIM, with known autoimmune genes in the top scJLIM and bulk JLIM results. More specifically, bulk JLIM yielded significance for 90% of the homogeneous (i.e., multi cell type) scJLIM hits compared to only 10% of the heterogeneous (cell-type-specific) ones (Supp. Figure 3a). A similar trend was found when applying bulk JLIM to cell subtypes. This test demonstrates that scJLIM is highly consistent with the pseudobulk methods when whole cell types have colocalization. It also underscores the shortcomings of pseudobulk methods, which have reduced power when colocalization does not cleanly fall into the predefined clusters.

We validated our results using a second PBMC dataset generated from 70 SLE patients and 70 healthy individuals^45^. We tested whether our scJLIM colocalizations would also be significant when tested within the same major cell types of the SLE dataset. In addition to retesting the OneK1K significant loci, we also retested a set of OneK1K non-significant loci as controls. We selected the same number of controls as positives for each trait-cell type combination since different colocalization rates might be expected in different cell types. We hypothesized that the OneK1K-significant loci would colocalize more often in the SLE dataset compared to the controls. Among loci that were significant only with scJLIM (and not bulk) in OneK1K, 15 of them (17%) replicated at p-value<0.01 in the SLE dataset (enrichment p-value = 1.8×10^-5^) (Figure 3e). Loci that were originally significant with both scJLIM and bulk JLIM replicated at a higher rate (60%, enrichment p-value = 5.4×10^-20^). This is to be expected, given that loci also identified with bulk JLIM tended to be stronger or to involve more cells. The imperfect replication rate is likely also a result of the nearly 10-fold reduction in sample size of the validation dataset and its potentially different cell states introduced from the SLE patients. Overall, this analysis reaffirms our confidence in the new colocalizations we observed with scJLIM.

To better understand the potential causes of missing regulation at GWAS loci, one recent study by Mostafavi et al. investigated the systematic differences in SNP and gene properties between bulk discovered eQTLs and GWAS loci^46^. This study emphasized the fact that bulk eQTLs tend to be located proximal to gene transcription start site locations, whereas GWAS variants tend to be relatively distal. Our results take those of previous study one step further showing that, at current sample sizes and power, we can only ascertain bulk colocalization for those few GWAS loci that are located proximally to the transcription start sites (TSS) of the eGenes they regulate (p-value=2.6×10^-5^) (Figure 3f). Conversely, the scJLIM-specific colocalized loci were further from the TSS of their respective eGenes (p-value=0.0067), while still being closer to them than background (p-value=0.015). Aligned with these observations, we also noted that the scJLIM-specific hits tended to have somewhat higher pLI scores (Supp. Figure 3b). From an evolutionary perspective, we speculate that it might be too deleterious for GWAS genes to be perturbed in all cells of a major cell type, especially if they are conserved genes. Therefore, selection might only allow most disease-causing variants to persist in the population if they alter regulatory elements of more specialized cell subtypes, producing the pattern we observe.

Amongst our scJLIM colocalizations was a significant (Padj = 0.02) finding for the *ETS2* gene in myeloid cells for the IBD trait. We investigated this finding further because a recent study validated the relevance of this gene in this context. That study identified that the IBD risk variant in this locus alters *ETS2* expression by disrupting binding of a transcription factor (TF) coded for by the *SPI1* gene^47^. Our analysis showed the non-classical monocytes (CD16+) to have the highest proportion of colocalizing cells, with additional colocalizing cells in the classical monocytes and conventional dendritic cell (cDC) populations (Supp. Figure 4a). We hypothesized that the *SPI1* TF would be more highly expressed in the colocalizing cells since the TF must be present for the causal SNP to dampen its regulatory effects. As expected, the *SPI1* TF had a similar pattern of expression across the cells (Supp. Figure 4b) with our colocalizing cells having significantly higher expression of this TF (Supp. Figure 4c). These findings thus serve as a validation that we detected the correct cell subpopulations for the colocalization of this gene with the IBD trait.

For most traits, we observed that within any major cell type, colocalizations tended to be cluster among the different predefined cell subclusters (Supplementary Tables 2). For instance, when analyzing the PBC trait and T cells (self-reactive T cells are known to infiltrate and target the bile ducts in PBC), we identified a subset of colocalizations biased towards naïve CD4+ T cells and others biased towards CD8+ T effector memory cells. Thus, we further clustered these scJLIM hits (Padj<0.10) by their per-cell colocalization significance patterns across cells without requiring the predefined subclusters. This was done by converting the per-cell p-values to Z-scores and calculating pairwise Pearson correlations between the loci. This enabled us to identify clusters of loci that potentially regulate their genes through the same T cell subtypes (Figure 3g). We also plotted uniform manifold approximation and projections (UMAPs) for these examples, indicating which cells contributed most to the global Cauchy significance (Figure 3h). One such cluster included the genes *ICAM3* and *TNFRSF14*, both of which had colocalizations primarily in the naive CD4+ T cells region. Previous studies have shown that these genes and the others have T cell functions related to the self-reactive role of T cells in this disease. For instance, functional experiments demonstrated that *ICAM3* is involved in early adhesion between CD4+ T cells and antigen presenting cells, which is an important precursor process for helper T cell stimulation^48^. Another study showed that *TNFRSF14* can serve as a co-stimulatory protein for naïve CD4+ T cells, and it can also regulate T-cell self-reactivity^49^. Knockout of *TNFRSF14* increased the susceptibility of mice to develop an autoimmune-like hepatitis (similar to the PBC trait) when stimulated with concanavalin A^50^. We further confirmed that these genes were not merely marker genes specific to the Naïve T cells (Supp. Figure 3c), exemplifying how regulatory variant effects are frequently orthogonal to expression levels at cell-type resolution. This demonstrates how scJLIM can help identify potential connections among GWAS genes, which has been challenging in the past. While only one example, our results for the other cell types and traits can also be analyzed in a similar manner to generate new inferences and hypotheses.

### scJLIM identifies cell subpopulations linked to neurological disease loci

We next investigated cell-type-specific colocalizations in common neurological diseases—an area where previous efforts have had limited success. We analyzed a large snRNA-seq dataset from postmortem prefrontal cortex brain samples of 378 aged individuals^51^, paired with nine GWAS datasets spanning highly prevalent neurodegenerative and neuropsychiatric diseases^52–60^. We applied scJLIM across eight major cell types: inhibitory neurons, *CUX2*+ (upper layer) excitatory neurons, *CUX2*-(deep-layer) excitatory neurons, microglia, astrocytes, endothelial cells, oligodendrocytes, and oligodendrocyte precursor cells (OPCs).

Compared to bulk tissue and subcluster-level JLIM, scJLIM identified a substantially higher number of colocalizations (Figure 4a). Neuronal subtypes, including inhibitory and excitatory neurons, yielded the greatest number of hits (Figure 4b), likely reflecting increased statistical power due to higher representation in the dataset. Importantly, scJLIM recapitulated several previously reported colocalizations, such as the rs4663105–*BIN1* locus in microglia for Alzheimer’s disease (AD), and rs6537401–*LSM6* in neurons and oligodendrocytes for ADHD, while also uncovering numerous novel associations, including rs2070902–*NDUFS2* in excitatory neurons in AD, rs10099100–*XKR6* in oligodendrocytes in ASD, and rs1401129–*UBE2E3* in astrocytes in bipolar disorder (BIP) (Supplementary Tables 3,4).

**Figure 4.**
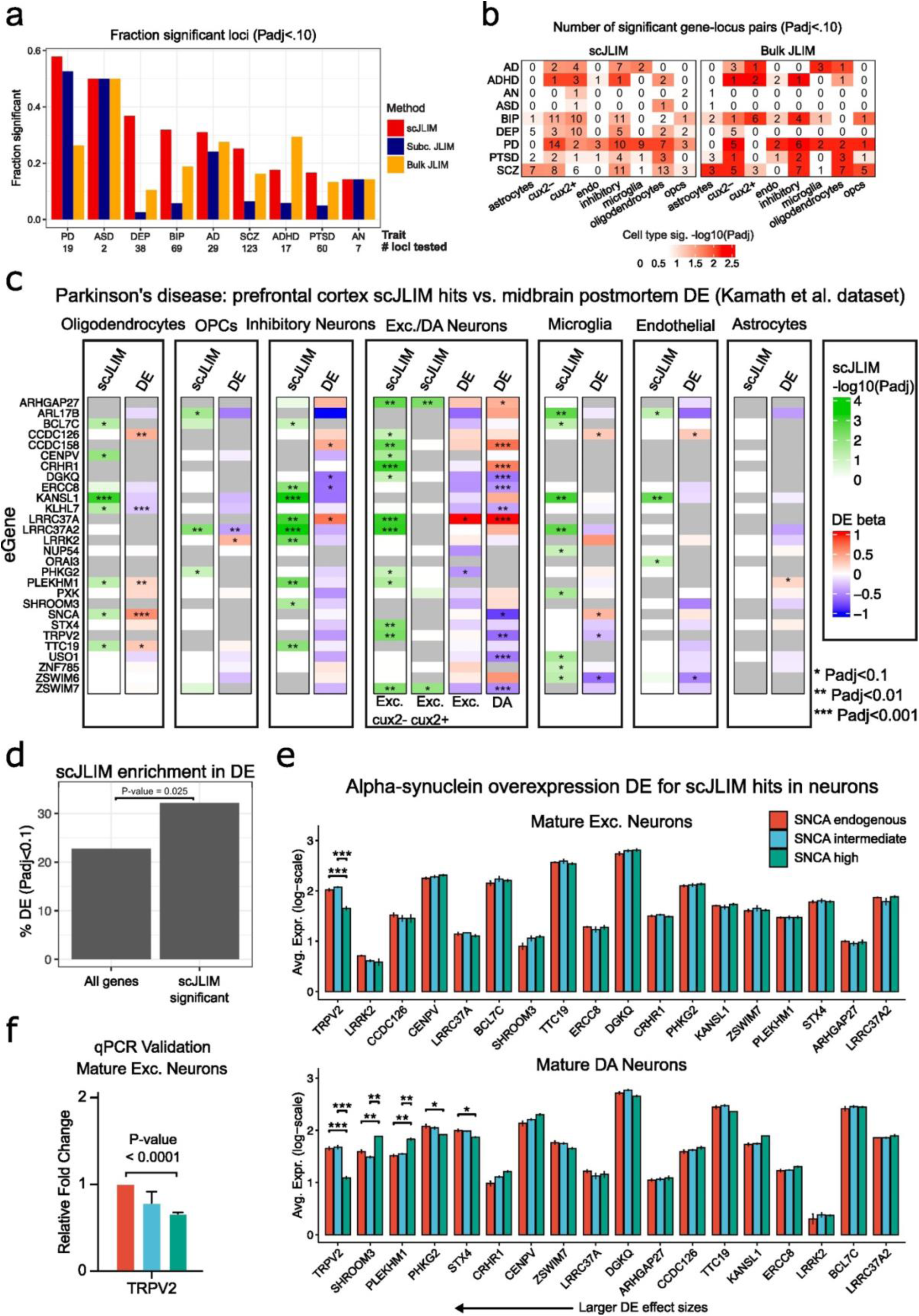
Application of scJLIM to the ROSMAP prefrontal cortex snRNA-seq dataset paired with nine GWAS datasets of neurodegenerative and neuropsychiatric diseases. a. Comparing the fraction of tested loci that are significant with either scJLIM, subcluster pseudobulk JLIM, or bulk JLIM. These results are aggregated over all eight major cell types. The total number of tested loci are reported under each set of bars for a trait. b. The number of significant gene-locus pairs for each trait in each cell type, shown separately for scJLIM (left) and bulk JLIM (right). Color indicates p-values from meta-analyzing all tests per trait-cell type pair with the Cauchy combination test. Here, cux2-denotes upper layer excitatory neurons, cux2-denotes deep-layer excitatory neurons, endo denotes endothelial cells, and opcs denote oligodendrocyte precursor cells. c. Overlap between scJLIM eGenes for Parkinson’s disease (PD) and significant DE genes between PD patients and healthy controls in a separate dataset of postmortem midbrain samples from Kamath et al.^57^. Gray color indicates that the gene was not tested in the indicated cell type. d. The percentage of scJLIM hits that are DE in the same cell types in the midbrain Kamath et al. dataset^57^ compared to background. The background included the shown gene-cell type combinations that were tested both in analyses. For excitatory neuron colocalizations, we included comparisons to either midbrain excitatory (Exc.) or dopaminergic (DA) neurons. The p-value was computed with a hypergeometric test for enrichment. e. Expression of scJLIM PD excitatory neuron hit eGenes in iPSC-derived cortical excitatory and DA neurons with pathologic brain-level alpha synuclein expression (*SNCA*-intermediate and *SNCA*-high) versus controls (*SNCA*-endogenous) at maturation. Only *TPRV2* shows significant downregulation in both cell types. f. Quantitative PCR validation of the *TRPV2* gene expression change in mature cortical neurons (e top). Expression is relative to that of *GAPDH*.

One disease where scJLIM particularly outperformed bulk colocalization was Parkinson’s disease (PD). We next investigated whether scJLIM colocalizations show disease-associated expression changes using an independent snRNA-seq dataset from postmortem midbrain samples from ten patients with PD or Lewy body dementia (DLB), and eight neurotypical controls^61^. Many scJLIM-significant genes were significantly differentially expressed (DE) between patients and controls in the same cell types (Figure 4c), with a significant enrichment for colocalized genes among DE genes (p = 0.025, Figure 4d). Interestingly, we observed strong concordance between colocalizations in deep-layer (*CUX2*-) excitatory neurons and DE genes in midbrain dopaminergic (DA) neurons, raising the possibility of shared regulatory mechanisms between these cell types— although the absence of DA neurons in our eQTL dataset and the limited presence of excitatory neurons in the midbrain snRNA-seq PD dataset precluded direct testing.

We further evaluated our scJLIM eGenes from neuronal cell types using a CRISPRi-induced synucleinopathy (CiS) model^62,63^ of α-synuclein (αS) toxicity, encompassing cortical excitatory and DA neurons. The CiS model includes clones with pathologic brain-level *SNCA* expression (*SNCA*-OE: *SNCA*-high and *SNCA*-intermediate) and a control clone (*SNCA*-endogenous). Cells were transduced at DIV0, with mature cortical excitatory neurons harvested at DIV28 and mature dopaminergic neurons at DIV60 for bulk RNA-seq (three replicates per clone). Several of the scJLIM neuron eGenes showed significant differential expression in *SNCA*-OE neurons compared to the *SNCA*-endogenous lines (Figure 4e). Notably, the direction of gene expression changes in *SNCA*-OE neurons aligned with the direction of effects of the corresponding GWAS risk variants on the expression of same genes. This observation suggests the presence of possible interactions or shared pathways between αS toxicity and our scJLIM hits.

Lastly, since the scJLIM eGene *TRPV2* (Transient Receptor Potential Cation Channel Subfamily V Member 2) was significantly downregulated in both cortical excitatory and DA *SNCA*-OE lines, we further validated its expression change by performing quantitative PCR (qPCR). This was done using the mature CiS-cortical excitatory neuron model (Figure 4f). Consistent with our previous bulk RNA-seq result, *TRPV2* had reduced gene expression for the cells with pathologic brain-level *SNCA* expression (*SNCA*-OE). *TRPV2* is broadly expressed across different brain cell types and is a key player in responding to diverse physical, chemical, and thermal stimuli^64^. While several other TRP family genes have previously been linked to PD^65^, the involvement of *TRPV2* in PD pathogenesis has not been explored in detail, marking this as a potentially novel insight.

## Discussion

In this work we described our approach to testing for eQTL-GWAS colocalization in scRNA-seq data by leveraging continuous representations of cell-states. Through simulations, we illustrated how bulk colocalization is susceptible to signal dilution or loss of power when merging signals from multiple distinct eQTL. We further showed how subclustering based colocalization is susceptible to clustering accuracy and resolution assumptions, all of which scJLIM were robust to. In applying our method to a PBMC dataset paired with autoimmune GWAS datasets, we uncovered roughly twice as many colocalizations as with bulk or subcluster pseudobulk eQTL mapping and showed that our new results were enriched for relevant genes. We validated our scJLIM-specific hits in a second PBMC dataset and observed concordance between our localized cell populations and those from experimental evidence or other methods. In general, each trait had multiple major cell types and subpopulations that were identified as relevant, and this is consistent with other studies which have similarly integrated scRNA-seq data with genetics^45,66^. By further demonstrating that scJLIM-specific hits were located further from transcriptional start sites, we provide evidence that cell subtype-specificity is key to resolving the missing heritability problem.

Application of scJLIM to a large snRNA-seq brain prefrontal cortex dataset also revealed new colocalizations with neurodegenerative and neuropsychiatric disease GWAS. Through an exploration of our PD results, we found significant overlap between our results and DE genes associated with PD pathology, both *in vivo* and *in vitro*. Notably, with scJLIM, we found a novel colocalization at the *TRPV2* locus mediated through the GWAS lead SNP, rs4566208, in vulnerable neuronal cell types in PD. This SNP had been found to colocalize with the gene, *ZSWIM7* at the brain tissue level in earlier studies^67^ but not *TRPV2*. We subsequently validated that *TRPV2* was significantly downregulated in DA neurons in postmortem brains of PD patients, as well as in iPSC-derived neurons with pathologic brain-level α-synuclein expression. This observation likely points towards direct and indirect pathogenic mechanisms mediated through *TRPV2*, a plasma membrane channel that plays a role in maintaining intracellular calcium (Ca^2+^) homeostasis^68^. The aggregation or overexpression of α-synuclein can impair the trafficking and expression of membrane proteins^69^, including *TRPV2*. Moreover, chronic cellular stress, inflammation, and the disruption of calcium homeostasis may further suppress *TRPV2* transcription or promote its degradation^70,71^. Together, these factors suggest that sustained downregulation of *TRPV2* may exacerbate neuronal vulnerability in PD. A recent murine model study has shown that *TRPV2* activation in the midbrain substantia nigra with induced degeneration regulates calcium influx, increases dopamine release, and improves locomotor activity in rats^72^. Thus, our findings around *TRPV2* open up possibilities for follow-up studies.

In AD, we noted several previously reported and novel colocalizations. Surprisingly, many of these new colocalizations were located in the inhibitory neurons. Since neurons cell types contain many cell subpopulations and ambiguous cluster boundaries, scJLIM may have substantially better power compared to bulk colocalization in such scenarios. Overall, these results highlight new genes and cellular contexts to be further characterized in AD and the other tested traits.

There are several limitations to our method and analyses conducted. The interaction-eQTL mapping step can be computationally burdensome in terms of runtime and memory usage, especially when applied to large scRNA-seq datasets. We took various measures to reduce this burden, but it will likely remain a limiting factor as scRNA-seq datasets continue to scale in size. Improvements in the computational efficiency will be necessary to increase statistical power through bigger sample sizes. Furthermore, as described in Nathan et al.^19^, linear mixed models can generate false positive interaction eQTLs if the eGene is both strongly differentially expressed (across the interaction) and very lowly expressed in one of the cell types. We reduced the chance of obtaining such false positives by restricting our tests to eGenes expressed above a minimum threshold, conducting our analyses within each major cell type separately, and by screening each gene-locus pair for inflated significance when permuting genotypes (Methods). While those authors suggest using a Poisson or negative binomial model, our simulations and genotype permutation tests (not shown) also pointed to excess false positives with this approach. This highlights the need for future work to understanding the sources of such false positives and improving the models accordingly. A third limitation of scJLIM, and colocalization testing in general, regards the interpretation of results as causative. We note that colocalization is required for but insufficient on its own to claim causality. For example, one might identify a colocalization in two different cell types, but it may be the case that the eGene only causally impacts the trait in one of the cell types or through an unmeasured cell type altogether. In spite of these limitations, scJLIM will help to expand our lists of disease gene candidates and their relevant cellular contexts for mechanistic follow-up studies.

More broadly, we tested whether the inclusion of the relevant context without the guesswork involved in assigning relevant cell states resolves the disconnect between the effects on gene expression and phenotypes. Our results suggest that taking cellular context into account is likely a part of the solution. However, even with the two-fold increase in the number of successful co-localizations, we are still far from explaining most of regulatory effects on complex traits.

## Methods

### scJLIM details

For the eQTL mapping step of scJLIM, we modeled the normalized, scaled, per-cell expression of the eGene as a function of genotype, various cell- and individual-level covariates (e.g., library size, age, sex, batch, etc.), each expression PC, and the interaction between genotype and each expression PC. Specifically, we use linear mixed models with a random effect over individuals, to account for the non-independence of different cells from the same individuals. The structure of one such model is shown below for including *K* PCs (equation 1).

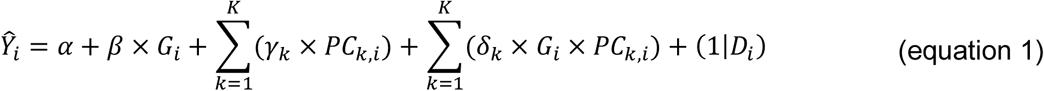

*Y*_*i*_ is the expression of the eGene in cell *i*. *G*_*i*_ is the SNP genotype of the donor that cell *i* is from. *PC*_*k*,*i*_ is the k^th^ PC and (1|*D*_*i*_) is the donor random effect. Additional covariates can be included for other individual- and cell-level variables, but these are omitted here for simplicity. The coefficients include *α* for the intercept, *β* for the persistent genotypic effect, *γ*_*k*_ for each PC effect, and *δ*_*k*_ for each genotype-by-PC interaction effect. We fit each model using the JuliaCall R package^73^ to access the increased computational performance of mixed-model fitting in Julia.

We then used each SNP’s fitted model to compute its total genotypic effect in each cell, conditional on the cell’s PC coordinates (i.e., the marginal effect). For a given cell, cell *i*, this is computed as in (equation 2), below.

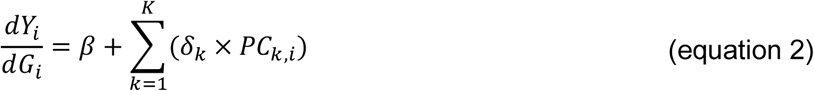

From this, we can see that the PC location of cell *i* determines whether its genotypic effect on expression is more or less than that of the persistent effect *β*. Then, we calculate the variance of this marginal genotypic effect with the delta method^74,75^, as in (equation 3), below.

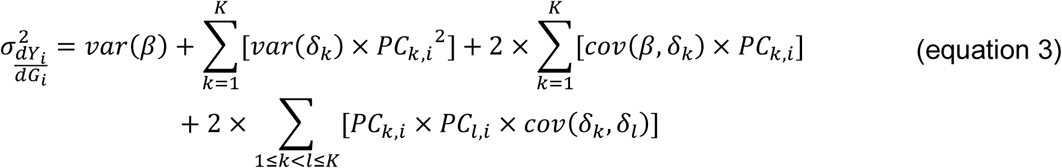

Here, we use *k* and *l* to refer to distinct PCs because we must include the covariance between multiple genotype-by-PC interaction term coefficients in the equation. We take the square root of this variance to get the standard error of the marginal genotypic effect in cell *i*. Then, we compute a Z-score for each SNP-cell pair by dividing the marginal effect by its standard error as 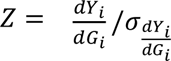 and calculate a two-sided p-value from this statistic with the cumulative distribution function of a standard normal. The p-values for all SNPs in a cell are then used as eQTL p-values for downstream colocalization testing per cell. We run the JLIM colocalization test as originally implemented. One difference, however, is that we only generate a single large null distribution (set to N=100,000 for this study) to compute the colocalization p-values in each cell, since it would be too computationally expensive to generate null distributions separately for each cell. This is possible because the null distribution only depends on the GWAS summary statistics, the eQTL sample size, and the reference LD matrix. We also reduced runtime by enabling scJLIM to run tests in parallel and share large data objects across nodes of a cluster, reducing RAM usage.

As noted in the main text, we take the JLIM p-values from all cells and apply the Cauchy combination test to calculate a global p-value indicating whether there is any significant colocalization among the tested cells. This p-value combination method is uniquely suited to our scenario as it allows p-values to be combined even when the p-values are from non-independent tests. This works because, under the null, the sum of Cauchy-transformed p-values with any arbitrary dependency structure approximates a standard Cauchy distribution. Before applying this test, however, we must apply a correction to our per-cell p-values because of a property specific to the JLIM test. Since the JLIM null distribution is generated with no eQTLs present, real loci harboring strong eQTLs out of LD with the sentinel GWAS SNP will receive overly conservative p-values. Therefore, in our application, cells might get p-values unrealistically close to 1 if they harbor other eQTLs in the locus. This is a problem when applying the Cauchy combination test because the presence of even a few “extremely null” appearing p-values will negate any real colocalization signals that may be present in different cells, resulting in a non-significant global p-value. We implemented a simple correction for this by replacing the p-values that are greater than 0.5 with those sampled from a random uniform distribution in the range of 0.5 to 1.0. Lastly, we also implemented an elimination-based procedure to select cells that contributed to a globally significant p-value. This procedure works by iteratively excluding the most significant cell and recomputing the global p-value until the global p-value rises above a user-defined threshold. Once surpassing the threshold, the final set of excluded cells is annotated as the colocalizing cell subpopulation

### Simulation design details

For the GWAS simulations, we generated genotypes for 10,000 individuals using Hapgen2^26^. For the eQTL dataset, we simulated an additional 50 individuals that are derived from the same population as the GWAS individuals (i.e., they have the same LD structure). Then, we selected 100 random loci, each 200kb wide, and picked a single SNP to be the causal GWAS variant in each locus. The causal SNPs were chosen to be relatively close to the middle of each locus, and we required it to have a high MAF of at least 0.2. Then, we simulated a quantitative phenotype per locus using the causal SNP and ran a linear model GWAS. For each locus, we also generated 5 simulated scRNA-seq datasets using the SplatPop R package^28^. This enabled us to program cell-type-specific eQTLs into the data using our simulated genotypes at each causal GWAS SNP. Each scRNA-seq simulation was made with 500 genes, 50 individuals, and 25 cells per individual per cell type (either two or four cell types). To create imbalanced cell proportion simulations (eQTL population in 30% or 70% of all cells), we kept the size of the smaller population at 25 cells per individual and increased the size of the other population accordingly. Likewise, the four-cell-type simulations were kept at balanced cell proportions with the same number of cells as before. We assigned each cell type to have its own set of marker genes upregulated, though we kept this signal relatively weak to help emphasize the difference in performance between scJLIM and subcluster JLIM. To make the simulated data more realistic, we added additional structured variation in the dataset by creating correlations among a small set of randomly selected genes. This also ensured that there would be multiple “real” PCs to be found that did not have genotype-interactions. For running scJLIM, we included two PCs for the two-cell-type simulations and five PCs for the four-cell-type simulations. When comparing simulations with perfect clustering versus inaccurate clustering, we computed clustering accuracy with a Jaccard-coefficient-based metric. Specifically, clustering accuracy was calculated by first computing the maximum Jaccard coefficients between each ground true cluster and each computed cluster. We then calculated the mean of these values over the two cell types to use as our statistic. For the example in Supp. Figure 2b, we selected tests with values less than 0.8 to illustrate how subcluster pseudobulk colocalization is highly sensitive to clustering accuracy.

### OneK1K processing details

We preprocessed the OneK1K genotype data similarly to the original publication^29^. We first converted the genotype matrix to Plink format^76^ and then filtered on a SNP genotyping rate of 0.03, individuals genotyping rate of 0.03, a Hardy-Weinberg p-value threshold of 1×10^-6^, MAF of 0.1, and genomic relatedness parameter of 0.125. These filtered data were input to the Michigan Imputation Server^77^ Minimac3 (hg19), followed by additional filtering to a MAF of 0.1, and R2<=0.8. For the scRNA-seq data, we first removed cells where the quality control metadata had values for the fail_QC column equal to “True”. Then, we for each major cell type, separately, we normalized the data by each cell’s library size, multiplied by a scale factor of 10,000, and applied a log transformation. Then, we selected the highly variable gene subset of the data with Seurat^78^, regressed out the effects of mitochondrial gene percentage and residual library size effects, scaled the data, and ran batch correction using ComBat^31^ with the “pool_number” variable as the batch. Next, PCA was run on the batch-corrected data, and we computed a UMAP using 20 PCs and a k-nearest-neighbor graph (k=10). We did not re-cluster the cells, using the previously annotated subclusters for analyses where relevant. The number of PCs to use per cell type in scJLIM was determined by comparing the PCA eigenvalues to 1.5 times the Marchenko-Pastur maximum, selecting those PCs with eigenvalues above this value. The result of this was 12 PCs for T cells, 7 PCs for NK cells, 10 PCs for B cells, and 10 PCs for Myeloid cells. For our linear mixed model eQTL mapping with scJLIM, we used non-batch corrected, normalized expression data as the response variable. In addition to the genotype, PC, genotype-PC-interaction, and donor random effect terms, we included covariates for age, sex, batch, library size, mitochondrial percentage, and 7 genotype PCs (to capture population structure within this European ancestry cohort). The genotype PCs were computed using Plink2^76^. For bulk and subcluster bulk eQTL mapping, we generated pseudobulk samples by summing counts per gene over all cells per cluster for an individual and normalized the same way as before. We used the same donor-level covariates for bulk eQTL mapping and additionally included the same number of PCs computed on the pseudobulk data. The subcluster colocalization results were meta-analyzed also using the Cauchy combination test to serve as a fair comparison to scJLIM. As this dataset represented samples from European ancestry, we used reference LD information from the 1000 genomes project European data subset when running JLIM^27^. Lastly, the enrichment statistics shown in Supplementary Tables 2 and 4 are calculated as 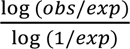, where *obs* is the observed fraction of cells in a subtype that have colocalization, and *exp* is the expected fraction based on the total number of colocalized cells.

### ROSMAP processing details

The ROSMAP scRNA-seq data was processed in a similar manner to the OneK1K dataset. We filtered out any cells labeled as either genotype doublets (i.e., from two individuals) or expression doublets. As in the original study, we exempted OPCs from removal of expression doublets which mislabeled the newly differentiated oligodendrocytes as doublets. Before computing PCs, we normalized the data, selected the highly variable gene subset of the data, regressed out the effects of library size and mitochondrial gene percentage, scaled the data, and applied batch correction (meta data column labelled “batch”). Next, PCA was run on the batch-corrected data, and we computed a UMAP using 30 PCs and a k-nearest-neighbor graph (k=10). We did not re-cluster the cells, instead using the previously annotated subclusters for analyses where relevant. For our linear mixed model eQTL mapping with scJLIM, we used non-batch corrected, normalized expression data as the response variable. We included the standard covariates for genotype, PCs, and genotype-PC interaction (10 PCs per cell type). We included additional covariates for age at death, sex, batch, postmortem interval, library size, mitochondrial percentage, and 3 genotype PCs (matching that of the original study). The genotype PCs were computed using Plink2^76^. Bulk eQTL mapping and colocalization was run the same way as with the OneK1K dataset, except that we could not include the batch variable because each individual had cells processed in multiple batches. However, as with the original study, we included bulk expression PCs to hopefully capture any residual batch effects. The 1000 genomes European ancestry reference LD panel was used when running JLIM^27^.

### SLE processing details

The SLE scRNA-seq data was processed in a similar manner to the OneK1K dataset. PCA was computed in the same manner as the OneK1K dataset with the same preprocessing steps and parameters. We subset the data to the individuals of European ancestry to match that of the GWAS data used. Prior to running PCA and UMAP, ComBat was run with the “batch_cov” meta data column to specify the batches of pooled individuals’ cells. For mixed model eQTL mapping, we included the standard covariates for genotype, PCs, and genotype-PC interaction (10 PCs per cell type). We included additional covariates for age, sex, batch, postmortem interval, library size, mitochondrial percentage, and 7 genotype PCs. Bulk eQTL mapping and colocalization was run the same way as with the OneK1K dataset. The 1000 genomes European ancestry reference LD panel was used when running JLIM^27^.

### GWAS processing and selecting loci

Minimal processing was conducted to the GWAS datasets. We removed structural variants and kept only SNPs on autosomes. We limited the SNPs to those we had genotype data for in the filtered OneK1K or ROSMAP datasets. The GWAS datasets selected were conducted in individuals of European ancestry to match that of the OneK1K or ROSMAP participants. For datasets with position information in hg38 coordinates, we lifted these over to hg19 coordinates using the common variant database from UCSC^79^. Then, we selected loci from each GWAS dataset separately, for testing with scJLIM. For a given GWAS, loci were defined by selecting all SNPs within 50kb of each genome-wide significant SNP (p-value < 5×10^-8^) that had a MAF over 0.1. Then, we merged loci that were intersecting. We excluded loci within chromosome 6 25mb-35mb to avoid the HLA regions. We then linked each locus to all protein-coding genes that were located within 500kb of the locus’ center.

### Single-cell transcriptomic analysis in post-mortem brain data

To validate our scJLIM-identified colocalizations in major central nervous system (CNS) cell types, we analyzed a publicly available snRNA-seq dataset from post-mortem midbrain samples of ten PD/DLB patients and eight healthy controls^61^. Cell type and subtype definitions from the original study were utilized, and snRNA-seq data were preprocessed according to the quality control recommendations provided in the original study. We performed differential expression analysis in all major cell types using MAST (model-based analysis of single-cell transcriptomes)^80^. We used a discrete hurdle model that assumes the snRNA-seq data follows a mixture of a binomial and normal distribution while accounting for pre-defined covariates. Sex, age, race, percentage of reads aligned to mitochondrial genes, and number of UMIs (log-scale) were included as fixed-effect covariates in the model. Disease status was used as the primary variable of interest. To account for dependencies between cells originating from the same individual, we included sample IDs as a random-effect covariate. The effect of disease status on gene expression was quantified using the beta value from the Wald test, where the sign of beta indicated the direction of the effect. We selected differentially expressed genes in each cell type as transcriptome-wide significant if we observed a Benjamini-Hochberg-adjusted p-value of less than 0.1.

### Human iPSCs culture, induced neuron differentiation, and CiS-model generation

To generate a neuronal model with pathologic brain-level alpha-synuclein (αS) expression (*SNCA-OE*), we transduced WTC11 human induced pluripotent stem cells (hiPSCs), which harbor a doxycycline-inducible-*NGN2* knocked-in at the *AAVS1* safe-harbor locus, with a lentiviral construct expressing *SNCA-IRES-mCherry*, as described in previous work^63^. A control group, with endogenous αS levels, was transduced with a lentiviral construct expressing *mCherry* alone. Following transduction, cells were sorted by flow cytometry (FACS) based on mCherry fluorescence intensity, which served as a proxy for *SNCA* expression, and replated at low density for single-cell clonal selection. Individual iPSC clones were expanded, karyotyped, and assessed for expression. Two clones from the *SNCA-OE* group and one from the control group were selected for further analysis and named *SNCA-high, SNCA-intermediate, SNCA-endogenous,* respectively. CiS-cortical neuron were differentiated and collected at maturation on DIV28^63^, and CiS-dopamine neuron were differentiated and collected at maturation on DIV 60 for total RNA extraction. For each clone, three biological replicates of differentiated neuronal samples were generated.

### Bulk RNA sequencing and analysis in CiS neurons

Total RNA was isolated from CiS-cortical and CiS-dopamine neurons using the PureLink™ RNA Mini Kit (Fisher Scientific; Cat. No. 12183018A), according to the manufacturer’s instructions.

For cDNA synthesis, RNA was reverse-transcribed using the SuperScript™ IV VILO™ Master Mix with ezDNase™ Enzyme (Fisher Scientific; Cat. No. 11766050). Paired-end strand-specific sequencing libraries were prepared from total RNA, and sequencing and annotation were conducted at Azenta Life Sciences.

Raw sequencing reads were mapped to the hg38/GRCh38 human reference genome, and transcript-level quantification based on GENCODE release 46 (GRCh38.p16) annotation was performed using Kallisto v0.46.2. We kept only protein-coding genes for subsequent analysis. All samples passed quality control checks for alignment rate, gene assignment rate, and mitochondrial mapping rates. We performed principal component analysis (PCA) and examined batch effects by inspecting the separation of samples by batch within the top principal components. Replicates were clustered by samples, not by batches, indicating consistency across batches. The replicates were included for quality control purposes and were not used in any additional analyses. All of our batch effect checks indicated consistency across batches.

For mature CiS-cortical and CiS-dopamine neurons, we performed differential expression analysis under three pairs of conditions: *SNCA-High vs SNCA-endogenous*, *SNCA-Intermediate vs SNCA-endogenous*, and *SNCA-High vs SNCA-Intermediate* using DESeq2 v1.42.1. Transcriptome-wide significant genes were selected with an absolute log2 fold change cutoff of 0.58 and a Bonferroni corrected p-value cutoff of 0.05.

### Quantitative PCR (qPCR)

Total RNA extraction and cDNA synthesis for DIV28 CiS cortical clones (*SNCA-OE* and *SNCA*-endogenous) were performed following the same steps as above. Quantitative PCR was performed using the PowerUp™ SYBR™ Green Master Mix (Fisher Scientific; Cat. No. A25777) with gene-specific primers. Reactions were performed on a QuantStudio Real-Time PCR System (Applied Biosystems QuantStudio 7 Flex) using the following thermal cycling conditions: 95°C for 15 seconds, followed by 40 cycles of 95°C for 1 second and 60°C for 20 seconds. Melt curve analysis was conducted to confirm amplification specificity. Relative gene expression was calculated using the 2^–ΔΔCt method, with GAPDH used as the internal reference gene.

## Data availability

The GWAS datasets were downloaded from various sources including the GWAS catalog^81^. From that database, we downloaded the following immune GWAS datasets with accession numbers in parentheses: PBC^35^ (GCST90061440), MS^34^ (GCST003566), eczema^38^ (GCST90244787), IGA^33^ (GCST003814), T1D^36^ (GCST90014023), RA^37^ (GCST90132223), and SLE^32^ (GCST003156). The IBD, Crohn’s, and UC GWAS datasets^39^ were downloaded from ibdgenetics.org from their files called EUR.IBD.gwas_info03_filtered.assoc, EUR.CD.gwas_info03_filtered.assoc, and EUR.UC.gwas_info03_filtered.assoc, respectively. The neurodegenerative and neuropsychiatric GWAS were downloaded from various sources. The bipolar (BIP) GWAS^60^ summary statistics from European individuals was downloaded from https://figshare.com/articles/dataset/bip2024/27216117?file=49760772. The AD GWAS^57^ was downloaded from https://vu.data.surfsara.nl/index.php/s/jVlyt1m9Bb2mAki. The ADHD GWAS^58^ was downloaded from https://figshare.com/articles/dataset/adhd2022/22564390. The AN GWAS^53^ was downloaded from https://figshare.com/articles/dataset/adhd2022/22564390. The ASD GWAS^56^ was downloaded from https://figshare.com/articles/dataset/adhd2022/22564390. The depression GWAS^54^ was downloaded from https://datashare.ed.ac.uk/handle/10283/3203. The PD GWAS^55^ was downloaded from the GWAS catalog with accession GCST009325. The PTSD GWAS^59^ was downloaded from https://figshare.com/articles/dataset/ptsd2024/26349322?file=47850151. The schizophrenia GWAS^52^ was downloaded from https://figshare.com/articles/dataset/scz2018clozuk/14681220?file=28198983. The Mayo clinic autoinflammatory gene panel was accessed at https://www.mayocliniclabs.com/test-catalog/overview/620092. The Priority Index database gene sets^43^ were downloaded from http://pi.well.ox.ac.uk:3010/. The ROSMAP scRNA-seq genotype data were accessed from Synapse under approved BWH study protocol 2025A003972. The OneK1K genotype matrix was downloaded from the Gene Expression Omnibus at accession GSE196829. The OneK1K cell annotations and quality control metadata^30^ were downloaded from https://github.com/immunogenomics/1K1K_QC.

## Code availability

The scJLIM tool is available as an R package on GitHub at https://github.com/j-mitchel/scJLIM. The code for generating our results in this manuscript is available at https://github.com/j-mitchel/scJLIM-Analysis.

---

**Supplementary Figure 1.**
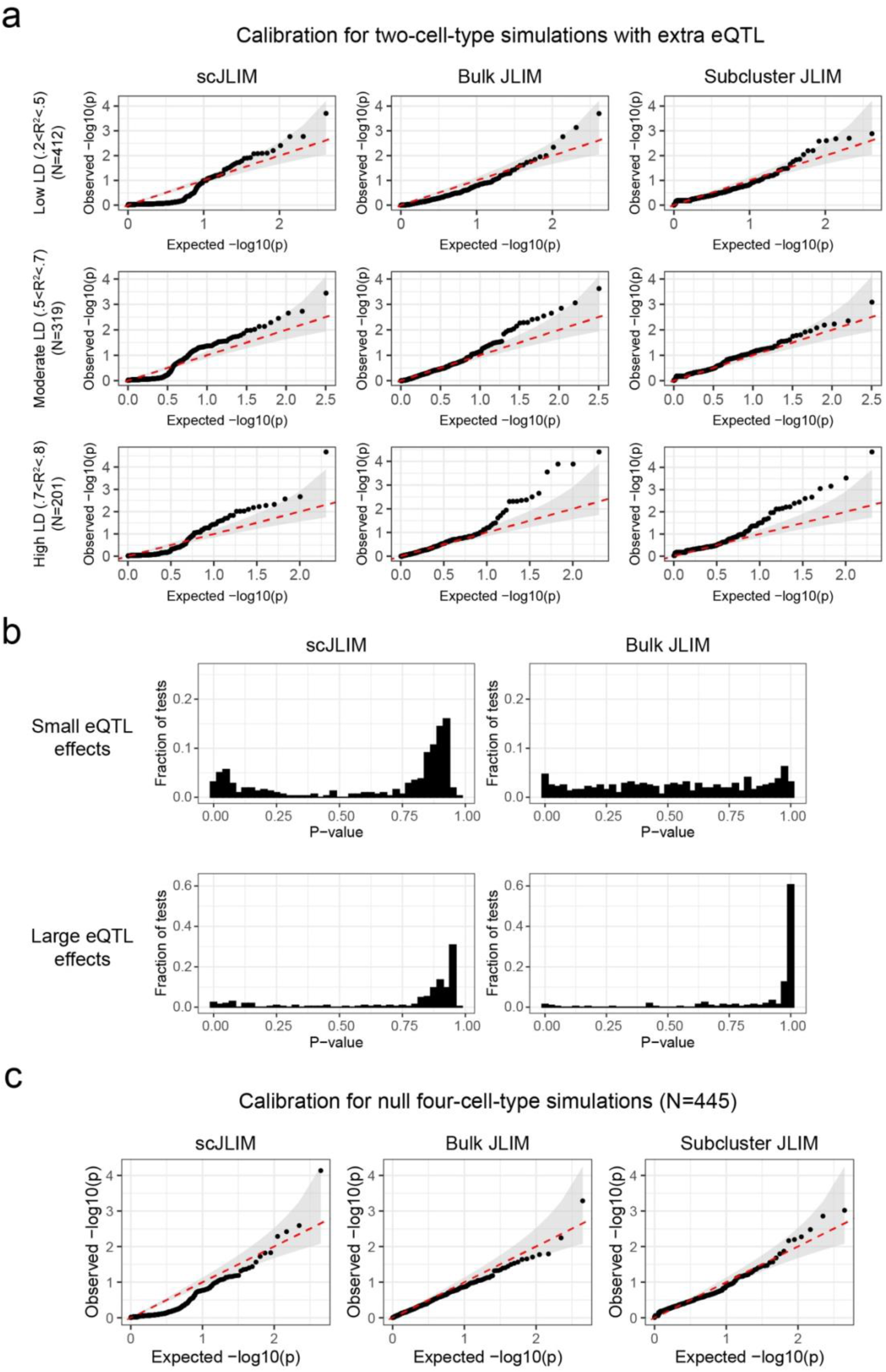
Additional simulation calibration results. a. Calibration results for simulations of the second null scenario in two cell types. These simulations contain one eQTL in one of the two cell populations, and this eQTL SNP is distinct from the causal GWAS SNP. The selected eQTL SNP was also chosen to be located within 50kb of the causal GWAS SNP and to have varying degrees of LD with it. The top, middle and bottom rows of plots have low, middle, and high LD values, respectively. These results are shown for each method separately in each column of plots. b. Comparing the scJLIM calibration p-value distributions for the eQTL effect sizes in (a) (top) versus larger effects (bottom). c. Calibration results for simulations of the first null scenario in four cell types.

**Supplementary Figure 2.**
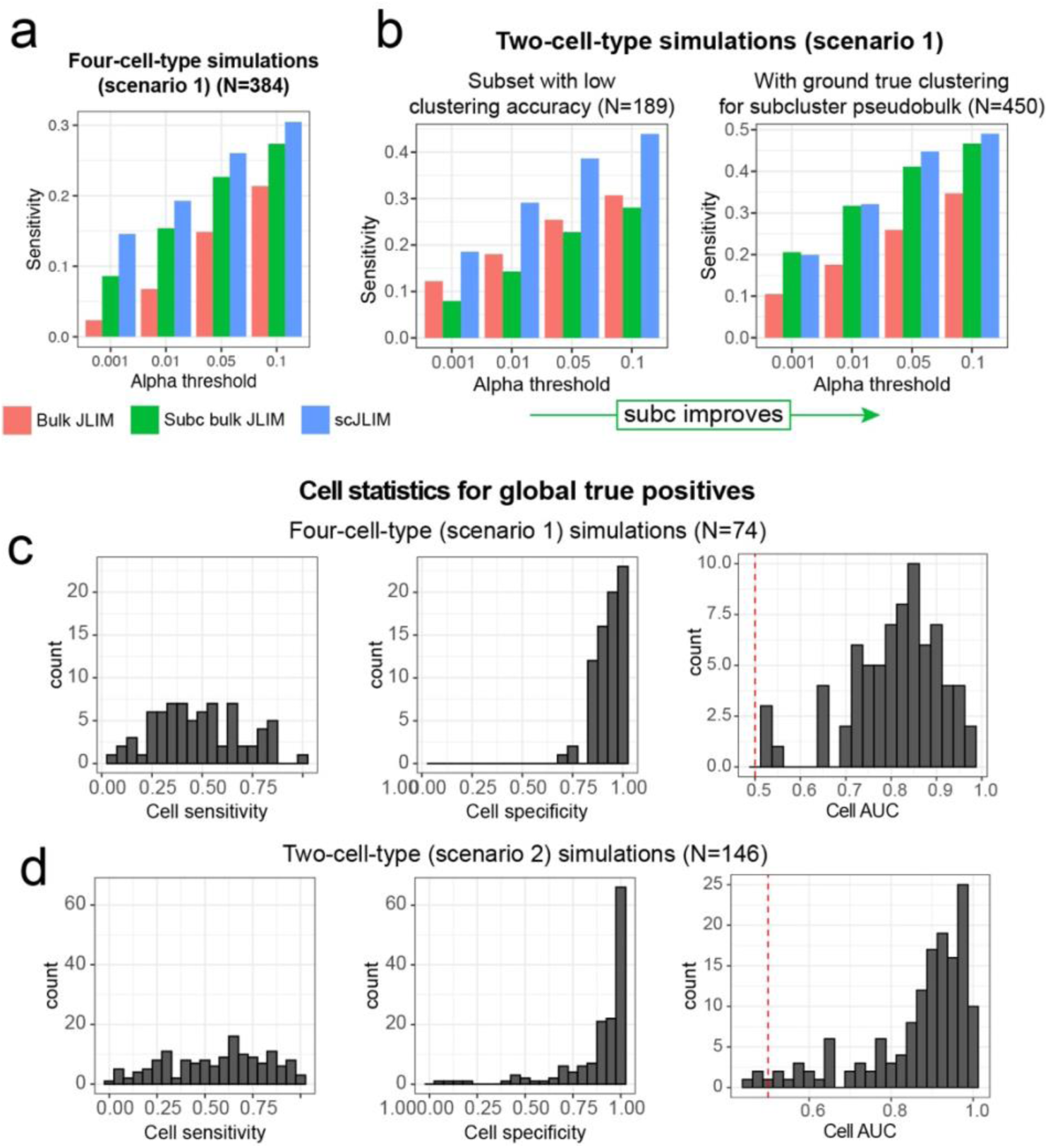
Additional results for the positive simulations. a. Sensitivity results for four-cell type simulations given positive scenario 1 (eQTL in one cell population, matching the GWAS SNP). b. Sensitivity results for the two-cell type positive simulations (scenario 1). The left plot shows those tests which specifically had low clustering accuracy based on a Jaccard coefficient metric comparing the real versus true clustering (Methods). On the right, we show the results for all tests when using the ground-true clustering labels for all cells. This illustrates how the performance of the subcluster pseudobulk colocalization matches that of our single-cell approach when we have perfect knowledge of each cell’s cluster assignments. c. Cell sensitivity, specificity, and ROC AUC for detecting the ground true cells that were given the eQTL for the same causal SNP as the GWAS data. This computed using the four-cell type, positive-scenario 1 simulations that yielded a global p-value less than 0.01. Significantly contributing cells were selected as those which pass our elimination-based procedure with the Cauchy combination test (Methods). d. As with (c), but for the two-cell-type positive scenario 2 simulations.

**Supplementary Figure 3.**
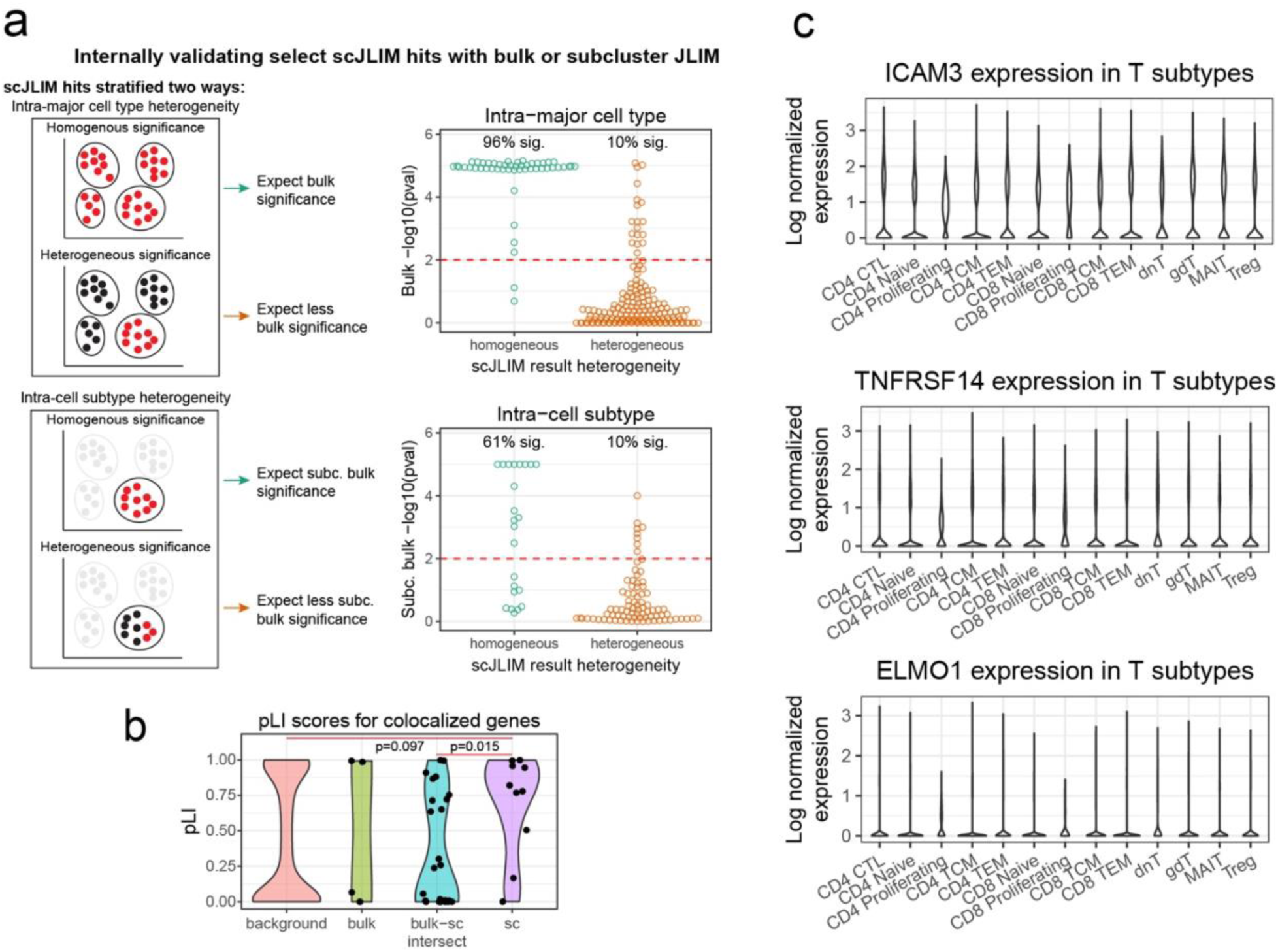
Additional analyses of the OneK1K dataset. a. Evaluating consistency between scJLIM and bulk or subcluster JLIM. We show bulk JLIM significance p-values for the scJLIM hits with low versus high intra-major cell type heterogeneity (right top). We also compared subcluster JLIM significance for scJLIM hits with low versus high intra-cell subtype heterogeneity (right bottom) (Methods). b. pLI scores are shown for the sets of eGenes identified in bulk JLIM only, scJLIM only, or the intersection of bulk and scJLIM. Significance was calculated using Wilcoxon rank-sum tests (two-sided). c. Gene expression levels in each annotated T cell subtype for the PBC trait genes colocalizing in Naïve CD4+ T cells. Expression is calculated as the log of the library size normalized UMI counts in each cell.

**Supplementary Figure 4.**
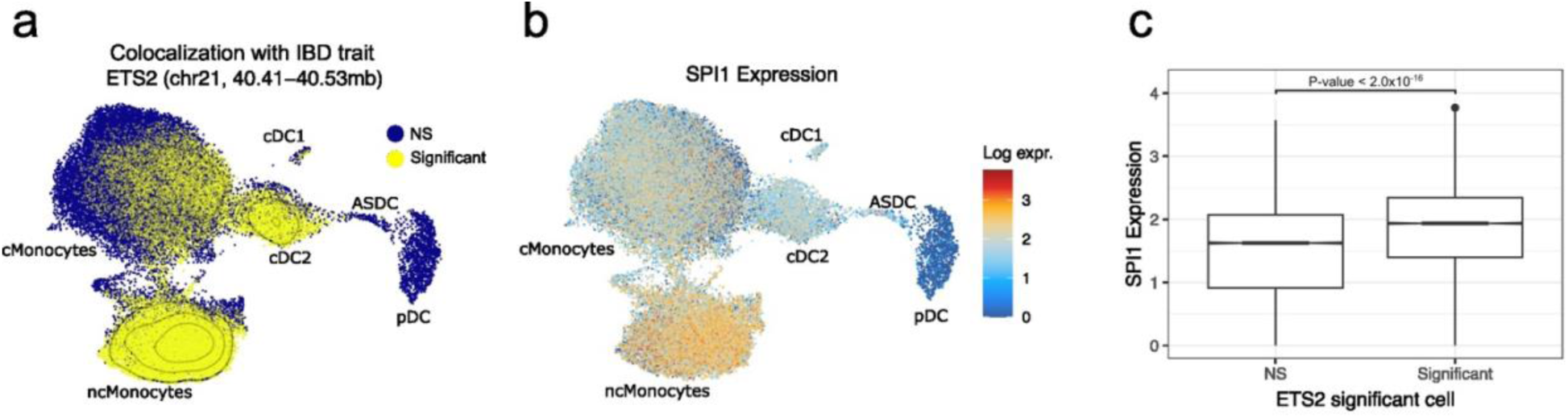
A validated IBD gene replicates in OneK1K dataset with scJLIM. a. scJLIM significantly contributing cells for a colocalization between *ETS2* eQTLs in myeloid cells and an IBD GWAS locus. b. Expression of the *SPI1* transcription factor in the myeloid cells. Expression values are log transformed. c. Expression of *SPI1* stratified by cells that contributed to the colocalization for *ETS2*.

**Supplementary Figure 5.**
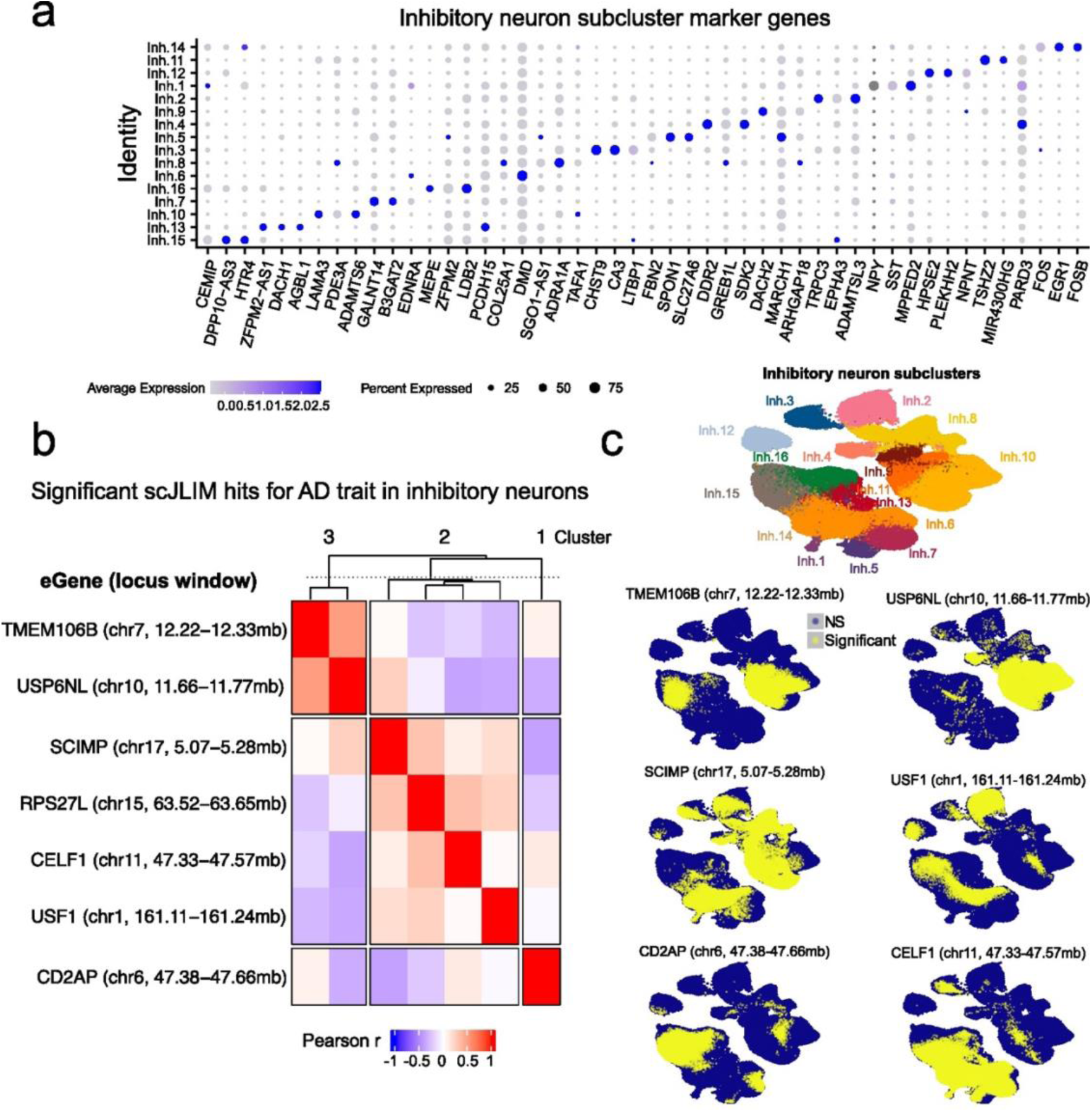
Additional analysis of AD loci colocalizing with inhibitory neurons in the ROSMAP dataset. a. Top marker genes differentially expressed between the inhibitory neuron subpopulations. b. Heatmap showing correlations between scJLIM significant (Padj<0.1) gene-locus pairs for the AD trait in inhibitory neurons. The correlations were calculated using the per-cell p-values in all inhibitory neurons, which were transformed to Z-scores prior to calculating each correlation. Pairs of results were clustered using hierarchical clustering and cut to select three clusters. c. Showing UMAP plots highlighting which cells contributed to global significance for select gene-locus pairs in (b). Significantly contributing cells were selected using the scJLIM per-cell p-values and include only those which pass our elimination-based procedure with the Cauchy combination test (Methods).

